# Auxin guides germ cell specification in *Arabidopsis* anthers

**DOI:** 10.1101/2020.10.13.337634

**Authors:** Yafeng Zheng, Donghui Wang, Sida Ye, Wenqian Chen, Guilan Li, Zhihong Xu, Shunong Bai, Feng Zhao

## Abstract

Germ cells (GCs) transmit genetic information from one generation to the next. Unlike animal GCs, plant GCs are induced post-embryonically, forming locally from somatic cells. This induction is coordinated with organogenesis and might be guided by positional cues. In angiosperms, male GCs initiate from the internal layers at the four corners of the anther primordia and are gradually enclosed by parietal cell (PC) layers, leading to a concentric GC-PC pattern.^1,2^ However, the underlying mechanism of GC initiation and GC-PC pattern formation is unclear. Auxin affects pattern formation^3^ and anther development.^4–11^ However, whether GC formation involves auxin remains unknown. We report that the auxin distribution in pre-meiotic anthers parallels GC initiation, forming a centripetal gradient between the outer primordial cells and the inner GCs. The auxin biosynthesis genes *TRYPTOPHAN AMINOTRANSFERASE OF ARABIDOPSIS 1* (*TAA1*) and *TRYPTOPHAN AMINOTRANSFERASE RELATED 2* (*TAR2*)^5,12^ are responsible for this patterning and essential for GC specification. *SPOROCYTELESS/NOZZLE* (*SPL/NZZ*, a determinant for GC specification)^13–15^ mediates the effect of auxin on GC specification, modulates auxin homeostasis, and maintains centripetal auxin patterning. Our results reveal that auxin is a key factor guiding GC specification in *Arabidopsis* anthers.

## Main

Germ cells (GCs) are initiated early in development in animals (during embryogenesis in ecdysozoa and chordata)^16^ and later in plants where they arise from somatic cells in the adult (after flowering in angiosperms).^1,17^ The mechanism for GC specification in plants is not well understood.^17–20^ In *Arabidopsis* and most other angiosperms, male GCs are derived from L2 cells at the four corners of the anther primordia. The primordial cells at the corners divide further to form outer parietal cells (PCs) and inner GCs (also termed archesporial cells, or ARs). GCs can be identified by their large cell size and big nuclei. Early pre-meiotic anther development occurs in five stages, as seen by changes to anther shape and cellular morphology (Fig. 1a).^2,21^ By stage 5, a concentric GC-PC pattern is fully established, germ cells (also termed pollen mother cells/PMC at this stage) are prepared for meiosis, and the anther exhibits a four-lobed butterfly-like shape (Fig. 1a). Many genes influence early anther development and form a complex gene regulatory network.^1,22,23^ In this network, the important upstream gene *SPOROCYTELESS/NOZZLE* (*SPL*/*NZZ*) is necessary and sufficient for GC initiation.^13–15^ However, the information about the positional signals guiding GC initiation pattern is very limited.

**Figure 1.**
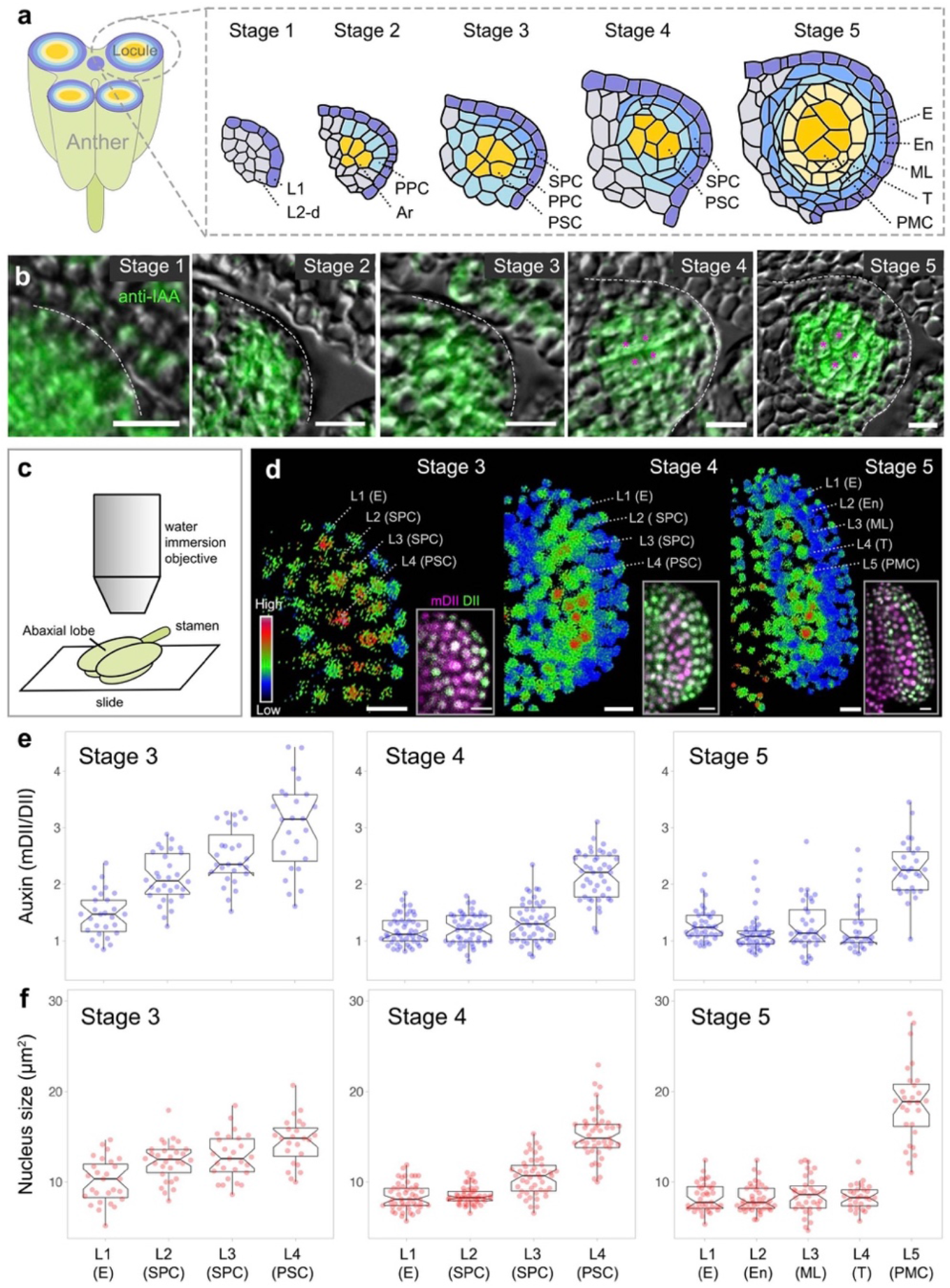
Dynamic auxin distribution patterns in wild-type pre-meiotic anthers in *Arabidopsis*. **a**, Schematic representation of microsporogenesis in early anther lobes. Left: Cross-section of a four-lobed anther. Right: The developmental stages of an anther lobe. Each cell lineage is marked by a specific color. L1, the outermost cell layer; L2-d, cells derived from the second layer; PPC, primary parietal cell; Ar, archesporial cell; SPC, secondary parietal cell; PSC, primary sporogenous cell; E, epidermis; En, endothecium; ML, middle layer; T, tapetum; PMC, pollen mother cell. **b**, Immunolocalization of IAA in cross-sections of anther lobes from stage 1 to stage 5. IAA gradually forms a centripetal gradient. The germ cells are marked with magenta stars. **c**, The method used for imaging R2D2 signals in abaxial anther lobes. **d**, Auxin signaling in anther lobes from stage 3–5, showing higher auxin levels (red) in germ cells. The signal intensity calculated from mDII/DII is displayed as a false color scale. The original images are shown in the insets. L1–L4 indicate the cell layers from outside to inside. The corresponding names of the cell layers are referenced in brackets. **e-f**, Quantification of auxin signaling input (mDII/DII) (**e**) and nucleus size (**f**) in different cell layers of stage 3–5 anther lobes. Note the association between auxin levels and germ cell specification (characterized by large nucleus size). The number of samples per layer are: L1=27, L2=30, L3=27, and L4=24 from four different stage-3 locules; L1=49, L2=48, L3=46, and L4=45 from six different stage-4 locules; L1=37, L2=41, L3=34, L4=28, and L5=28 from four different stage-5 locules. The individual data points are colored and plotted on the boxplot. The box indicates the interquartile range (IQR), the whiskers show the range of values that are within 1.5*IQR and a horizontal line indicates the median. The notches represent the 95% confidence interval for each median. Scale bars represent 10 μm.

Auxin is a morphogen-like compound that plays a pivotal role in pattern formation in plant morphogenesis,^3^ including anther development.^6–9^ Auxin is essential for anther initiation,^4^ pollen development, anther dehiscence, and other aspects of post-meiotic anther development,^7–9,24^ but its function in microsporogenesis, i.e. germ cell initiation and specification during pre-meiotic anther development, is unknown. Therefore, we investigated the role of auxin in guiding GC specification.

We first studied auxin distribution during early anther development using immunostaining and auxin bio-sensors. Using antibodies specific to the major form of auxin, indole 3-acetic acid (IAA), we labeled native auxin in early anthers from stage 1 to stage 5 (Fig. 1b, Fig. S1). In stage 1 anthers, auxin was homogeneously distributed in all cells. In stage 2, auxin spread into lateral domains. Starting from stage 3, auxin gradually congregated in central GC domains, accompanying the differentiation of the cell layers; stage 5 anthers displayed a clear difference in auxin levels between GCs and PCs (Fig. 1b, Fig. S1).

R2D2 is a ratiometric auxin bio-sensor that is used to indicate the relative *in vivo* auxin concentration at the cellular level.^25,26^ We mapped R2D2 signals in abaxial anther lobes (Fig. 1c-f). The longitudinal view of anther lobes yielded a fine map of auxin distribution patterns from stage 3 to stage 5: in stage 3, the auxin gradient was steep, with a minimum in the outer cells and a maximum in the inner cells; from stage 4 to stage 5, auxin was concentrated in the cells of the GC domains (Fig. 1d-e), which are characterized by their large nuclei (Fig. 1f).

The auxin signals detected through immunolocalization and bio-sensors are the combined output of auxin transport,^27^ metabolism (including biosynthesis,^28^ modification,^29^ and degradation^30^), and complex signaling pathways that stimulate downstream events.^31^ To pinpoint the particular components responsible for the centripetal auxin distribution and its potential effects on GC specification, we conducted RNA-seq and genetic manipulation experiments. RNA-seq analysis revealed a variable expression pattern for auxin-related genes in stage 3 and stage 4 anthers, but no clear hints to pinpoint whether auxin polar transport, metabolism, or signaling pathways potentially play a key role in early anther development (Fig. S2). We then created transgenic plants that interfered with a range of auxin-related functions in early anther development (Fig. S3) and used Alexander staining to evaluate GC formation.^20,32^ Phenotyping of these plants showed that altered auxin synthesis caused severe pollen defects (Fig. S3). These genetic data suggested that the regulations in auxin biosynthesis pathway may be important for microsporogenesis. Therefore, we examined the role of auxin biosynthesis in GC formation.

The L-Trp-dependent indole-3-pyruvate biosynthesis pathway is the main route for IAA production.^28^ This pathway includes enzymes encoded by genes in the *TRYPTOPHAN AMINOTRANSFERASE OF ARABIDOPSIS 1* (*TAA1*) and *YUCCA* (*YUC*) families. Because the *TAA1* family works upstream of the *YUC* family,^33^ and *TAA1* and *TRYPTOPHAN AMINOTRANSFERASE RELATED 2* (*TAR2*) are expressed in early anthers in our RNA-seq data, we focused only on *TAA1* and its close homolog, *TAR2* in our investigation of the role of auxin biosynthesis in GC formation. *TAA1* and *TAR2* were initially expressed throughout anther primordia, but became concentrated in GC domains by stage 5 (Fig. S4). In *wei8-1 tar2-2* (a weak allele of the *taa1 tar2* mutant^5^), around 40% of anthers formed pollen in two locules, ~20% harbored pollen in only one locule, and the remaining 40% were completely sterile (Fig. S5a). By contrast, the anthers of *wei8-1 tar2-1* (a strong allele of *taa1 tar2* ^5^) were completely devoid of pollen (Fig. S5a). Thus, TAA1 and TAR2 affect GC formation.

To investigate the role of auxin biosynthesis in centripetal auxin patterning and GC differentiation, we crossed R2D2 with *wei8-1/+ tar2*. In the F_3_ generation, as expected, auxin production was significantly reduced in *wei8-1 tar2-2* and *wei8-1 tar2-1* anthers (Fig. 2a-b), and auxin-responsive genes were broadly downregulated (Fig. S5b). Even though a weak centripetal auxin gradient persisted in *wei8-1 tar2-2* abaxial anther lobes, this gradient was lost in *wei8-1 tar2-1* (Fig. 2a-b). Consistent with the perturbation of the auxin distribution pattern, the nucleus size also changed in *taa1 tar2* mutants. In wild-type stage 3/4 anthers, the nuclei of GCs located in L4 were bigger than those of somatic cells (Fig. 1f), similar to those in *wei8-1/+ tar2-2* (Fig. 2c, Fig. S5a). However, in *wei8-1 tar2-2*, the L4 nuclei were significantly smaller (Fig. 2c) and *wei8-1 tar2-1* anthers had smaller nuclei in L4 and L3 cells (Fig. 2c).

**Figure 2.**
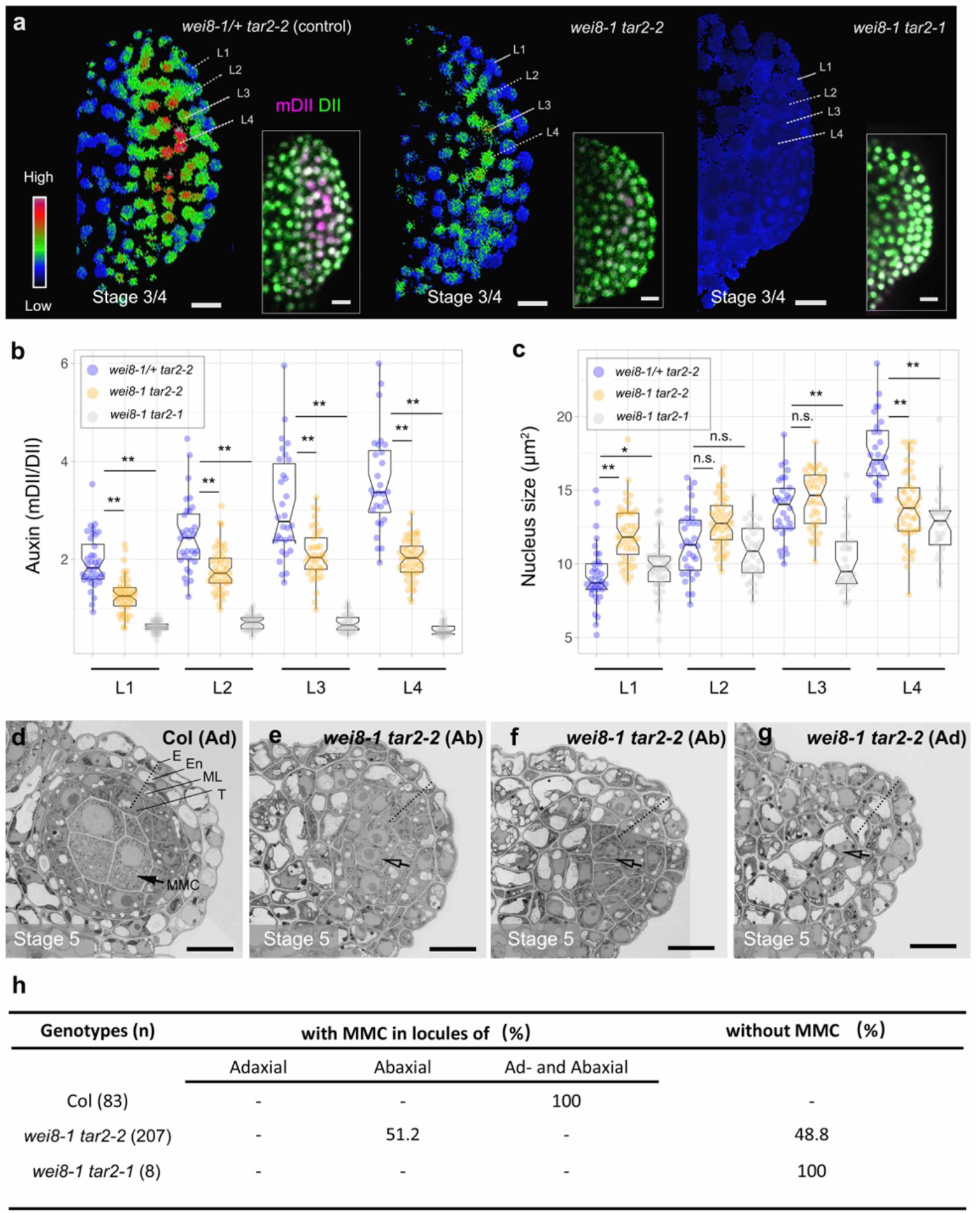
TAA1/TAR2 is indispensable for germ cell specification. **a**, Ratiometric image of R2D2 shows auxin signaling input in *taa1 tar2* anther lobes at stage 3/4. The signal intensity calculated from mDII/DII is displayed as a false color scale. The original images are shown in the insets. Compared with *wei8-1/+ tar2-2*, the auxin level is lower in *wei8-1 tar2-2* and extremely low in *wei8-1 tar2-1*. **b-c**, Quantification of auxin signaling input (**b**) and nucleus size (**c**) in stage 3/4 *taa1 tar2* anther lobes. As the auxin level decreases in the *taa1 tar2* mutants, the nucleus becomes smaller in the inner cells (L3 and L4), implying the loss of germinal cell fate. The number of samples per layer was: L1=36, L2=32, L3=30, and L4=28 from four control (*wei8-1/+ tar2-2*) anther lobes; L1=46, L2=47, L3=37, and L4=49 from five *wei8-1 tar2-2* anther lobes; L1=47, L2=35, L3=30, and L4=28 from four *wei8-1 tar2-1* anther lobes. n.s., no significant difference; * *p*<0.05; ** *p*<0.001 by Student’s *t*-test. **d-g**, Electron microscope images showing the transverse sections of wild-type (Col) adaxial (Ad) (**d**), *wei8-1 tar2-2* abaxial (Ab) (**e-f**), and *wei8-1 tar2-2* Ad locules (**g**). In (**d**), the normal MMC is marked with a black arrow, and different somatic cell layers are indicated along the dashed line. The impaired MMC (hollow arrow) and somatic cell layer (dashed lines) are marked in (**e-g**). Note that most cells in the *wei8-2 tar2-2* adaxial locules are vacuolated (**g**). **h**, Quantification of MMC formation in cross-sections of *taa1 tar2* anthers. Scale bars represent 10 μm.

In examination of the differentiation status of GC cells using histological analysis in stage 5 *wei8-1 tar2-2* anthers (Fig. 2d-g), we found in some of the abaxial locules, GC-like cells were visible, but the cell number and size were reduced (Fig. 2e). In other abaxial locules, cells were mostly vacuolated, which complicated the observation of GCs or well-differentiated PCs (Fig. 2f). All cells in the adaxial locules were vacuolated, so we were unable to identify GCs or well-organized PC layers (Fig. 2g). We quantified these phenotypes: 48.8% of *wei8-1 tar2-2* anthers failed to form GCs or well-differentiated PCs and 51.2% of the anthers only showed GCs at abaxial locules (Fig. 2h, Fig. S5c). The strong allele *wei8 tar2-1* contains only locules with vacuolated cells, which caused sterility (Fig. 2h, Fig. S5c). Thus, we concluded that *TAA1* and *TAR2* are required for dynamic auxin distribution during early anther development, and indispensable for GC specification.

How is auxin involved in GC specification? In *Arabidopsis*, SPL/NZZ is the determinant for GC differentiation,^13–15^ and the *SPL/NZZ* expression pattern overlaps with the distribution of auxin (Fig. S6a).^20^ In *spl/nzz* anthers, ARs (early stage of GCs) and their adjacent cells stop differentiating at stage 3.^14,15^ In stage 4, cells become vacuolated, hindering the formation of GCs and well-organized PCs. To test if auxin affects GC specification by regulating *SPL/NZZ* expression, we probed *SPL*/*NZZ* for transcripts in *wei8-1 tar2-2* anthers by *in situ* hybridization. In 50% of anthers, *SPL*/*NZZ* transcripts were diminished or absent in the adaxial lobes, and completely absent in the rest (Fig 3a, Fig. S6a). We quantified *SPL*/*NZZ* transcripts using quantitative polymerase chain reaction (qPCR) in *taa1 tar2* flowers. *wei8-1 tar2-2* contained 70% fewer *SPL*/*NZZ* transcripts and *wei8-1 tar2-1* contained almost no *SPL*/*NZZ* transcripts when compared with wild type (Fig. 3b). To test whether auxin mediates *SPL/NZZ* expression at the transcriptional or at the post-transcriptional level, we examined *SPL/NZZ* promoter activity in *taa1 tar2* by crossing *pSPL:GUS* with *taa1/+ tar2*. In the F_3_ progeny, GUS signals became weaker in *wei8-1 tar2-2* and were absent in *wei8-1 tar2-1* anthers (Fig. 3C, Fig. S6b). Thus, auxin regulates *SPL/NZZ* expression at the transcriptional level.

**Figure 3.**
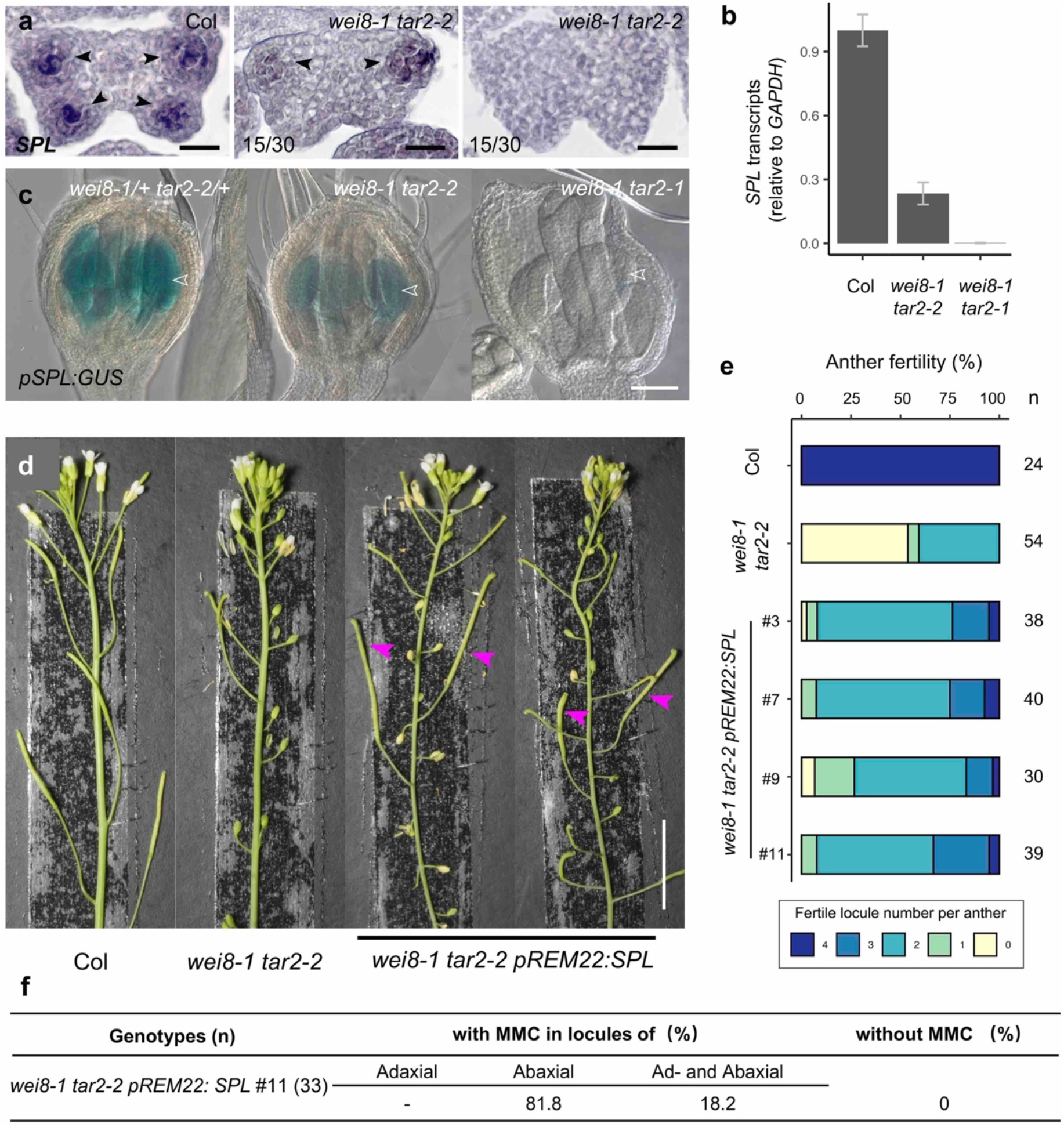
TAA1 and TAR2 activate SPL/NZZ transcription. **a**, *SPL/NZZ* transcripts in cross-section of stage-4 Col and *wei8-1 tar2-2* anthers, revealed by *in situ* hybridization. The *SPL*/*NZZ* signals are indicated by arrowheads. **b**, Quantification of *SPL/NZZ* transcripts in Col and *taa1 tar2* mutants by qPCR. Bars indicate the mean, and the error bars indicate standard error (*n* = 3 technical repeats). Three independent experiments yielded similar results. **c**, The *SPL* promoter activity shown by *pSPL:GUS* signals in Col and *taa1 tar2*. *pSPL* activity is reduced in *wei8-1 tar2-2*, and absent in *wei8-1 tar2-1* anthers (arrowheads). **d**, The fertility of *wei8-1 tar2-2* is partially restored by *pREM22:SPL*. Arrowheads (magenta) indicate fertile long siliques. **e**. Quantification of anther fertility in Col, *wei8-1 tar2-2,* and *wei8-1 tar2-2 pREM22:SPL* by Alexander staining assay. **f**. Quantification of MMC formation in cross-sections of *wei8-1 tar2-2 pREM22:SPL* anthers. Scale bars represent 20 μm in (**a**), 1 mm in (**c**), and 1 cm in (**d**).

To test if auxin affects GC specification by regulating *SPL/NZZ* transcription, we ectopically expressed *SPL/NZZ* under another AR-active promoter, *pREM22*.^34^ If this transgene complemented the *taa1 tar2* phenotype, that would indicate *SPL/NZZ* is downstream of auxin in mediating GC specification. As expected, *pREM22:SPL-Myc* partially restored the fertility of *wei8-1 tar2-2* (Fig 3d-e).^34^ In a *wei8-1 tar2-2 pREM22:SPL-Myc* plant, GCs appeared in adaxial locules of 18.2% of the anthers, which were fully sterile in *wei8-1 tar2-2* (Fig. 3f, Fig. 2h). The proportion of abaxial locules with GCs increased from 51.2% in *wei8-1 tar2-2* (Fig. 2h) to 81.8% in *wei8-1 tar2-2 pREM22:SPL-Myc* (Fig. 3f). These data demonstrated that *SPL/NZZ* functioned downstream of auxin to mediate GC specification. On the other hand, as we found abolishment of a centripetal auxin gradient in *spl* mutant anthers (Fig. 4a-c), together with the reported suppression of *YUC* genes by ectopic expression of *SPL/NZZ*^35^, we conclude that *SPL/NZZ* also has a feedback regulatory effect on the auxin biosynthesis pathway to maintain auxin homeostasis and distribution.

**Figure 4.**
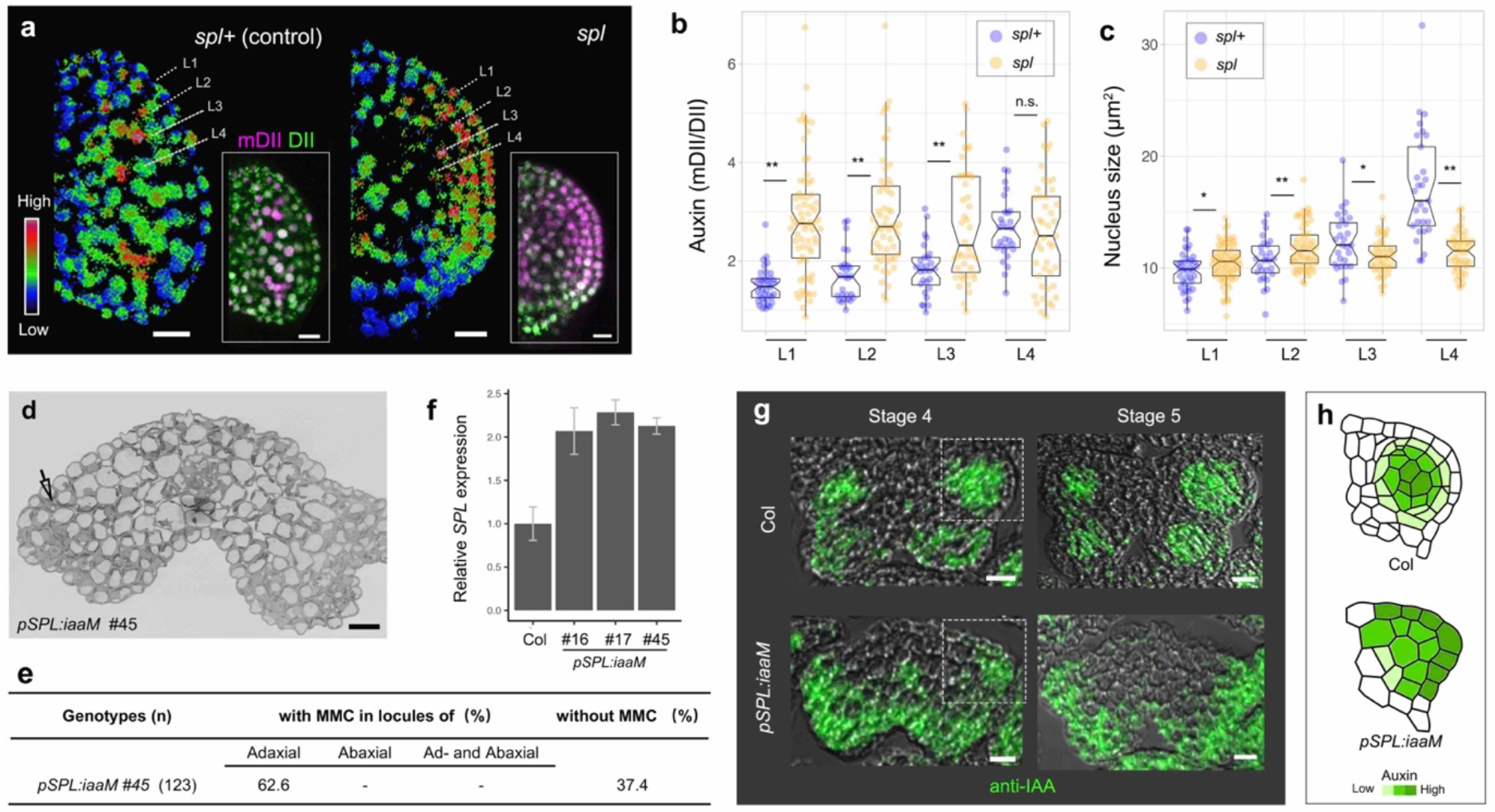
Auxin-SPL feedback loop for GC specification. **a**, Ratiometric image of R2D2 showing an increase of auxin in the outer cell layers of *spl* stage 3/4 anther lobes. The signal intensity calculated from mDII/DII is displayed as a false color scale. The original images are shown in the insets. **b-c**, Quantification of auxin signaling input (**b**) and nucleus size (**c**) in stage 3/4 *spl+*(control) and *spl* anther lobes. The auxin level was significantly increased in L1-L3 of *spl* anther lobes (**b**), while the nucleus size was significantly reduced in *spl* anthers, corresponding to the loss of GCs (**c**). The number of samples per layer was: L1=38, L2=27, L3=28, and L4=29 from four control (*spl+*) anther lobes; L1=68, L2=56, L3=42, and L4=45 from six *spl* anther lobes. n.s., no significant difference; * p<0.05; ** p<0.001 by Student’s *t*-test. **d**, Electron microscope images showing the transverse sections of a stage 5 *pSPL:iaaM* anther. All cells in the locules were vacuolated (arrowhead) and no GCs or somatic layers were distinguishable. **e**, Quantification of MMC formation in *pSPL:iaaM* anthers. **f**. qPCR showing *SPL* transcript levels in flowers from three different T5 *pSPL:iaaM* alleles and in controls. Numbers indicate alleles. Bars indicates the mean, and the error bars indicates standard error (*n* = 3 technical repeats). Three independent experiments yielded similar results. **g**, IAA immunolocalization in Col and *pSPL:iaaM* anthers at stage 4 and 5. IAA is distributed in the outer layers in *pSPL:iaaM*, similar to that in (a-b). **h**, Schematic representation of IAA distribution shown in (g, frame zones). Scale bars represent 10 μm.

Auxin deficiency results in a defect in GC specification; excess auxin also affects GC formation (Fig. S3, Fig. S7a-b). We quantified the abnormality of anther development in four independent T2 lines of *pSPL:iaaM* and three independent T_1_ lines of *pREM22:iaaM* (Fig. S7a). Around 20–60% of anthers in these transgenics were completely sterile and only 3.65% of the anthers were normal (Fig. S7a). Cells that normally differentiate into GCs in wild type did not differentiate, but became vacuolated (Fig. 4d). This deficiency occurred in 62.6% of abaxial anther lobes and the remaining 37.4% of anthers were completely sterile (Fig. 4d-e, Fig. S7c). In *pSPL:iaaM* transgenic lines, *SPL/NZZ* gene expression increased (Fig. 4f), consistent with upregulation by auxin (Fig. 3a-c). However, the centripetal auxin distribution pattern was severely disrupted (Fig. 4g-h). This abnormal auxin distribution pattern is similar to what we observed in the *spl* mutant (Fig. 4a-b). Given that *SPL/NZZ* negatively regulates auxin biosynthesis to maintain auxin homeostasis and patterning (Fig. 4),^35^ our observations in *pSPL:iaaM* anthers underline the importance of auxin control in GC specification.

In most higher animals, the GCs differentiate in early embryogenesis to form a germline carried by the soma. In contrast, in most land plants, GCs that are committed into meiosis are segregated from somatic cells during late development. In many angiosperms, such as *Arabidopsis*, male GCs are initiated at the four corners of anther primordia, but little was known about the positional information that induces GC initiation. Our data demonstrates that auxin plays a key role; in particular, auxin biosynthesis is indispensable. Through spatiotemporal interactions with the GC-determining *SPL/NZZ*, a centripetal auxin gradient forms and GCs are specified in the inner layers of anther lobes. When the auxin gradient is abolished, GC specification halts. The dynamic interactions between auxin and *SPL/NZZ* provide a new context for the spatiotemporal model of GC initiation and specification. This leads to several intriguing questions, such as how the homogenous auxin distribution is disrupted during anther development (possibly via polar auxin transport)?^6^ What is the detailed mechanism of auxin/SPL feedback? How do GCs differentiate and induce the centripetal auxin distribution? Answering these questions will shed further light on how GCs are induced and specified from somatic cells during plant development. In addition, this knowledge may improve our understanding of how plants and animals evolved such dramatically different routes for GC induction and differentiation.

## Methods

### Plant material and growth conditions

The *Arabidopsis* reporter lines (R2D2,^25^ *pSPL::GUS*,^20^ and *pSPL:SPL-myc*^20^) and mutants (*spl*,^14^ *wei8-1 tar2-1* ^5^, and *wei8-1 tar2-2*^5^) have been described previously. The seeds were sterilized, placed on Murashige and Skoog (MS) medium for germination, and cultured *in vitro*. After two weeks, the seedlings were transplanted to soil and grown under long-day conditions (16-h light/8-h dark; light bulb, Philips 28 W 840 neon, 4000 K, 103 lm/W) at 22°C.

### Plasmid construction and plant transformation

To overexpress auxin-related genes in anthers, the full-length cDNA was cloned and fused into an *SPL* promoter cassette.^20^ For generating RNAi lines, amiRNAs were designed following the instructions on the website (http://wmd3.weigelworld.org/cgi-bin/webapp.cgi); these were cloned into the *SPL* promoter cassette.^20^ These constructs were transformed into Col-0 using the floral dip method.^36^ T1 plants were screened on Murashige and Skoog (MS) medium containing glufosinate-ammonium; resistant plants were selected for further analysis.

The *REM22* promoter was chosen based on its activity in GC precursors (ARs).^34^ A 1022-bp region upstream of the start codon was cloned into the pEGAD and pCambia1305.1 target vectors to get *REM22* promoter cassettes; these were named pEGAD-REM22 and pCambia1305.1-REM22, respectively. To obtain the *pREM22:iaaM* construct, the *iaaM* coding sequence was cloned into pEGAD-REM22. It was then transformed into Col plants and transgenics were screened using glufosinate-ammonium *in vitro*.

The *pREM22:SPL-Myc* construct was generated by fusing the *SPL-Myc* coding sequence, which was cloned from the *pSPL:SPL-Myc* construct,^20^ into the pCambia1305.1-REM22 vector. *pREM22:SPL-Myc* was then transformed into *wei8-1/+ tar2-2* plants. *pREM22:SPL-Myc* transgenics were screened using hygromycin B and identified by PCR in T_2_ progeny. The primers used for cloning are listed in Table S1.

### Immunofluorescence localization assay

To detect IAA distribution, inflorescences were soaked in 3% (w/v) 1-ethyl-3-(3-dimethylaminopropyl) carbodiimide (Sigma Chemical, St. Louis, MO, USA, E6383) with 0.05% (v/v) Triton X-100, vacuum infiltrated for 1 h, and incubated in darkness at 4°C for 1 h. The inflorescences were rinsed three times in phosphate buffer (0.2 M, pH 7.4) for 10 min each, transferred to fixative (4% (w/v) paraformaldehyde in 0.2 M phosphate buffer and 0.1% (v/v) Triton X-100), vacuum infiltrated for 1 h, and incubated overnight at 4°C. Paraffin sectioning was performed as described previously,^37^ with a minor modification. The 7-μm sections were spread on poly-Lys-coated slides. After dewaxing, the slides were incubated in fixative again, and washed twice with washing buffer (0.2 M phosphate buffer, 0.1% (v/v) Tween 20) for 10 min each. For blocking, the sections were soaked in 10 mM phosphate buffer containing 3% (w/v) BSA blocking solution for 1 h (at room temperature) or overnight (at 4°C). The anti-IAA antibodies (Phytodetek, catalog no. PDM 09346/0096; diluted 1:150 in blocking solution) were added and the sections incubated for 3–4 h (at room temperature, 25°C) or overnight (at 4°C). After incubation, the samples were washed twice in 10 mM phosphate buffer containing 2.9% (w/v) NaCl, 0.1% (v/v) Tween 20, and 0.1% (v/v) BSA for 10 min at a time, and then washed with 10 mM PBS, 0.88% (w/v) NaCl, 0.1% (v/v) Tween 20, and 0.8% (w/v) BSA for 10 min. Anti-rabbit Alexa Fluor 594 (affinity anti-Myc antibody) and anti-mouse Fluor 488 (affinity anti-IAA antibody) secondary antibodies were diluted 1:500 in blocking solution and incubated for 4 h at room temperature. Two washes (15 min each) with 10 mM PBS, 0.88% (w/v) NaCl, 0.1% (v/v) Tween 20, and 0.8% (w/v) BSA were followed by a 1 min rinse in 10 mM PBS. Images were photographed using a Zeiss microscope (Axio Imager D2), processed by ZEN lite 2011 (blue edition; Carl Zeiss), and edited with Photoshop CS6 (Adobe Systems). At least three biological replicates were performed and similar results were obtained in all experiments.

### Confocal microscopy

Flowers between stages 7 and 9 were dissected and the sepals were removed. The anthers (at stages 3–5) were excised from the meristem, and quickly placed on a slide with double-sided tape (3M) to hold the abaxial locules upward. A drop of water was placed on the anthers and they were imaged using an LSM 710 confocal microscope (Carl Zeiss, Oberkochen, Germany) equipped with a water immersion objective (W Plan-Apochromat 20x/1.0 DIC VIS-IR). Fluorescence images were analyzed using ZEN LITE 2011 software (black edition, Carl Zeiss). *wei8-1 tar2-2* and *spl* were sterile and in different genetic backgrounds than R2D2. To compare the R2D2 signal in different genetic backgrounds, we chose *wei8/+ tar2-2* and *spl/+* (which are as fertile as wild type; see supplemental data) as the controls for *wei8 tar2* and *spl*, respectively.

To quantify the R2D2 signals, contours of nuclei were manually selected in the mDII channel using the elliptical selection tool in Fiji freeware, and regions of interest (ROIs) were added to the ROI manager. The area of the nuclei and the mean gray values of different ROIs in mDII and DII channels were measured. The mDII/DII ratio was calculated in Microsoft Excel. The results were plotted using the PlotsOfData web tool.^38^

To generate ratiometric images of R2D2, ratios between signal intensities of each pixel from the mDII and DII channel were calculated using Fiji; signal intensities in both channels below 10–60 (based on the average signal intensities between the nuclei in the mDII channel) were set to 0 in ratio images to subtract the background.

### RNA-sequencing

Stage 3 and stage 4 stamens were dissected and pooled separately for each replicate. RNA was extracted using RNeasy Plant Mini Kit (Qiagen, Cat NO. 74904) according to the manufacturer’s instructions. Libraries were constructed using the TruSeq Stranded Total RNA Sample Prep Kit (Illumina). The RNA-seq libraries were sequenced using a HiSeq2000 Pair End 2×100bp at the Peking University BIOPIC High-throughput Sequencing Center. The original image data generated by the sequencing machine were converted into sequence data via base calling (Illumina pipeline CASAVA v1.8.2). Then subjected to standard quality control (QC) criteria to remove all of the reads that fit any of the following parameters: (1) reads that aligned to adaptors or primers with no more than two mismatches, (2) reads with >10% unknown bases (N bases), and (3) reads with >50% of low-quality bases (quality value #5) in one read. Finally, 1308.3 Gb (94.4%) of filtered reads were left for further analysis after QC, and reads mapped to rRNA were discarded. After that, the remaining reads were mapped to the TIR9 reference genome using Bowtie 2 and TopHap. The RNA-seq data are deposited at Beijing Institute of Genomics Data Center (https://bigd.big.ac.cn/?lang=en) (BioProject: PRJCA003607).

### Alexander staining assay

To stain pollen, flowers were dissected from the inflorescence, opened using tweezers, and soaked in Alexander staining solution^32^ for at least 10 h. The stamens were dissected from the flowers, placed on slides, and sealed with chloral hydrate solution (4 g chloral hydrate, 1 mL glycerol, and 2 mL deionized water). To compare the different genotypes, flowers were dissected at the same developmental stage.

### *In situ* hybridization

*In situ* hybridization was performed according to a previously described protocol.^39^ Images were taken using a Zeiss microscope (Axio Imager D2). Primers used for preparing the probes targeting *TAA1*, *TAR2*, and *SPL* transcripts are listed in table S1.

### Histological analysis

Cross-sections of anthers were obtained by paraffin sectioning. The procedures were conducted according to Wang et al., 2010.^37^ Inflorescences were collected at the same developmental stage in order to compare the phenotypes between different lines. Anthers at stages 4–11 were chosen to quantify locule sterility. Quantitative analysis was carried out using Microsoft Excel.

For electron microscopy, flowers from stages 7–9 were selected from the inflorescences and soaked in 4% (w/v) paraformaldehyde and 2.5% (w/v) glutaraldehyde (Sigma, G5882). After vacuum infiltration for 1 h, the samples were incubated at 4°C overnight. They were post-fixed in 2% (w/v) OsO_4_ (Ted Pella, 18451) in phosphate buffer (0.1 M, pH 7.4) at room temperature for 90 min. The staining buffer was replaced with 2.5% (w/v) ferrocyanide (Sigma, 234125) in phosphate buffer (0.1 M, pH 7.4) at room temperature for 90 min. After being washed three times in 0.1 M phosphate buffer, the samples were incubated with filtered thiocarbohydrazide (Sigma, 223220) at 40°C for 45 min. Then the samples were fixed in unbuffered 2% OsO_4_ for 90 min, followed by incubation in 1% (w/v) uranyl acetate (Zhongjingkeyi, China) aqueous solution at 4°C overnight. Then, they were incubated in a lead aspartate solution (0.033 g lead nitrate (Sigma, 228621) in 5 mL 0.03 M aspartic acid (Sigma, 11189, pH 5.0)) at 50°C for 120 min, and dehydrated through a graded ethanol series (50%, 70%, 80%, 90%, and 100% ethanol, 10 min each) and pure acetone. The samples were embedded in Epon-812 resin (SPI, 02660-AB). Ultrathin sections (50 nm) were cut with a diamond knife (Diatome, MC16425) and imaged using a scanning electron microscope (Zeiss Gemini 300) with a resolution of 3 nm/pixel and dwell time of 2–5 μs.

### Quantitative Real-Time PCR

Total RNA was extracted from *Arabidopsis* inflorescences using E.Z.N.A Plant RNA kit (OMEGA, R6827-01). RNA samples were digested using RQ1 RNase-free DNase (Promega, M6101). For first-strand cDNA synthesis, 1 μg of RNA was used with ReverTra Ace qPCR RT Kit (TOYOBO, Code No.FSQ-101). The SYBR Premix Ex Taq (TaKaRa, RR420A) was used to carry out quantitative real-time PCR using the Applied Biosystems 7500 real-time PCR system. *GAPDH* was used as internal reference. The sequences of primers used are listed in Supplemental Table S1.

### *β*-Glucuronidase (GUS) staining assay

For GUS staining, whole *pSPL:GUS* inflorescences were soaked in X-Gluc solution (100 mM sodium phosphate buffer, 10 mM EDTA, 0.5 mM K_3_Fe(CN)6, 3 mM K_4_Fe(CN)6, 0.1% (v/v) Triton X-100) containing 1 mg/mL of *β*-glucuronidase substrate X-gluc (5-bromo-4-chloro-3-indolylglucuronide)), and vacuum infiltrated for 1 h in darkness. The inflorescences were incubated at 37°C in darkness for 16 h. After staining, the samples were dehydrated in a graded ethanol series [70%, 85%, 95%, and 100% twice (v/v)] until the chlorophyll was completely removed. Flowers were then dissected and placed on the slide, sealed with chloral hydrate solution, and photographed using a Zeiss microscope (Axio Imager D2). Images were processed using ZEN lite 2011 (blue edition; Carl Zeiss).

## Acknowledgements

We would like to thank Jan Traas for critical reading of the manuscript, Weicai Yang, Dolf Weijers, Jose M. Alonso, and ABRC for providing the seeds. We also thank Yang Xu, Yun Zhang, and Fuchou Tang for the supporting on RNA-sequencing. We thank Rui Chen for advice about RNA-seq data analysis. We also thank Fanbo Meng for helping with plant breeding and sectioning of anthers. We thank the Core Facilities of Life Sciences and National Center for Protein Sciences at Peking University in Beijing, China, for assistance with confocal microscopy and TEM. We would be grateful to Yiqun Liu and Hongmei Zhang for their help of making EM sample and taking images. We also thank Linlin Li and colleagues (Institute of Automation, Chinese Academy of Sciences) for the assistance with electron microscopy (Zeiss Gemini 300) and their technical support. This work was supported by grants from the Ministry of Agriculture of the People’s Republic of China (Grant No. 2016ZX08009003-003, CARS-01-06, and 2016ZX08010001) and the National Natural Science Foundation of China (Grant No. 31630006) to S.N.,B.

## Author contributions

F.Z. and S.N.B. conceived the project. Y.F.Z, F.Z., and S.N.B. designed experiments, and wrote the manuscript. Y.F.Z, F.Z., and D.H.W. performed the experiment and analyzed the data. S.D.Y. contributed to the IAA immunolocalization assay. G.L.L performed RNA-seq analysis. W.Q.C and Z.H.X contributed to the experimental design and edited the manuscript. All authors read and approved the final manuscript.

## Additional information

Correspondence and requests for materials should be addressed to F.Z. or S.N.B.

---

**Figure S1.**
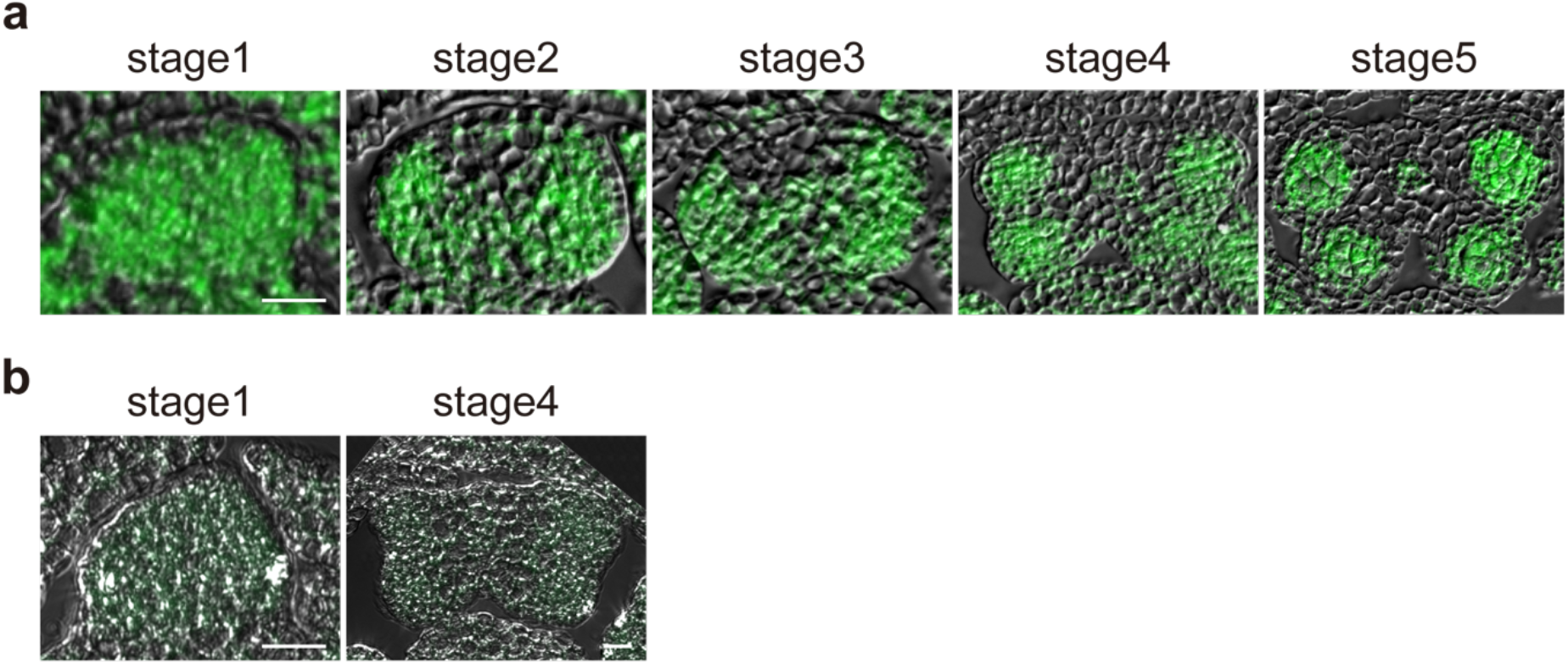
Dynamic IAA distribution during early anther development revealed by immunolocalization. **a**. Representative picture of IAA immunolocalization showing the dynamic IAA distribution pattern in cross-sections of Col anthers. Similar results were obtained from at least 3 replicates. **b**. Negative control of IAA immunolocalization on stage1 and stage4 Col anthers, without adding anti-IAA antibodies. Scale bars represent 10 μm.

**Figure S2.**
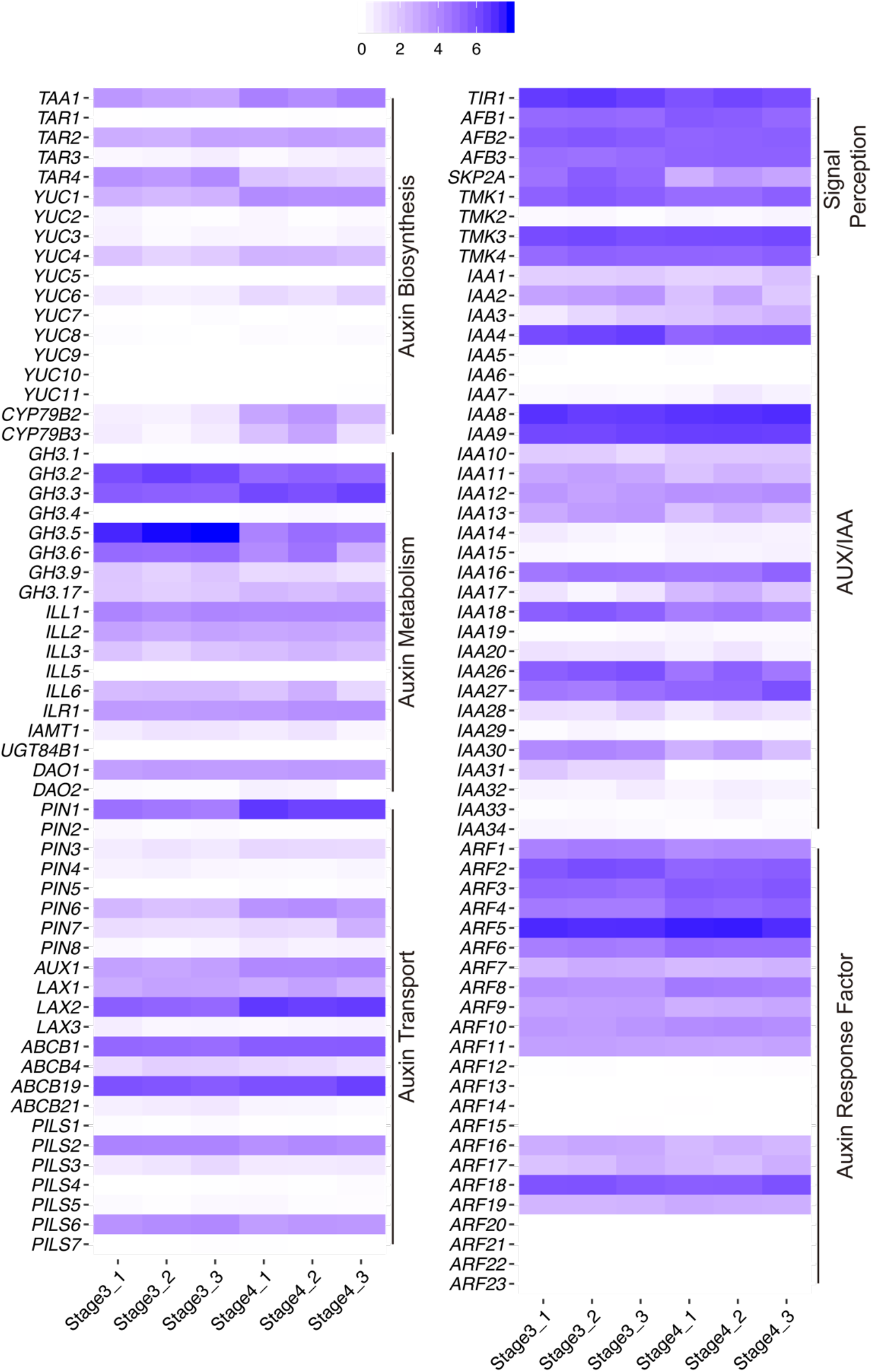
The expression profiles of auxin-related genes in Col stage 3 and 4 anthers. Heat map of auxin-related genes expression in stage 3 and stage 4 anthers by RNA-seq analysis. The color relatively represented log2(genes’ relative expression value plus 1). Stage3(4) _1, stage3(4) _2, stage3(4) _3 represented 3 replicates of RNA-seq results of stage 3(4) anthers. Almost all auxin pathways were activated in stage 3 and stage 4 anthers.

**Figure S3.**
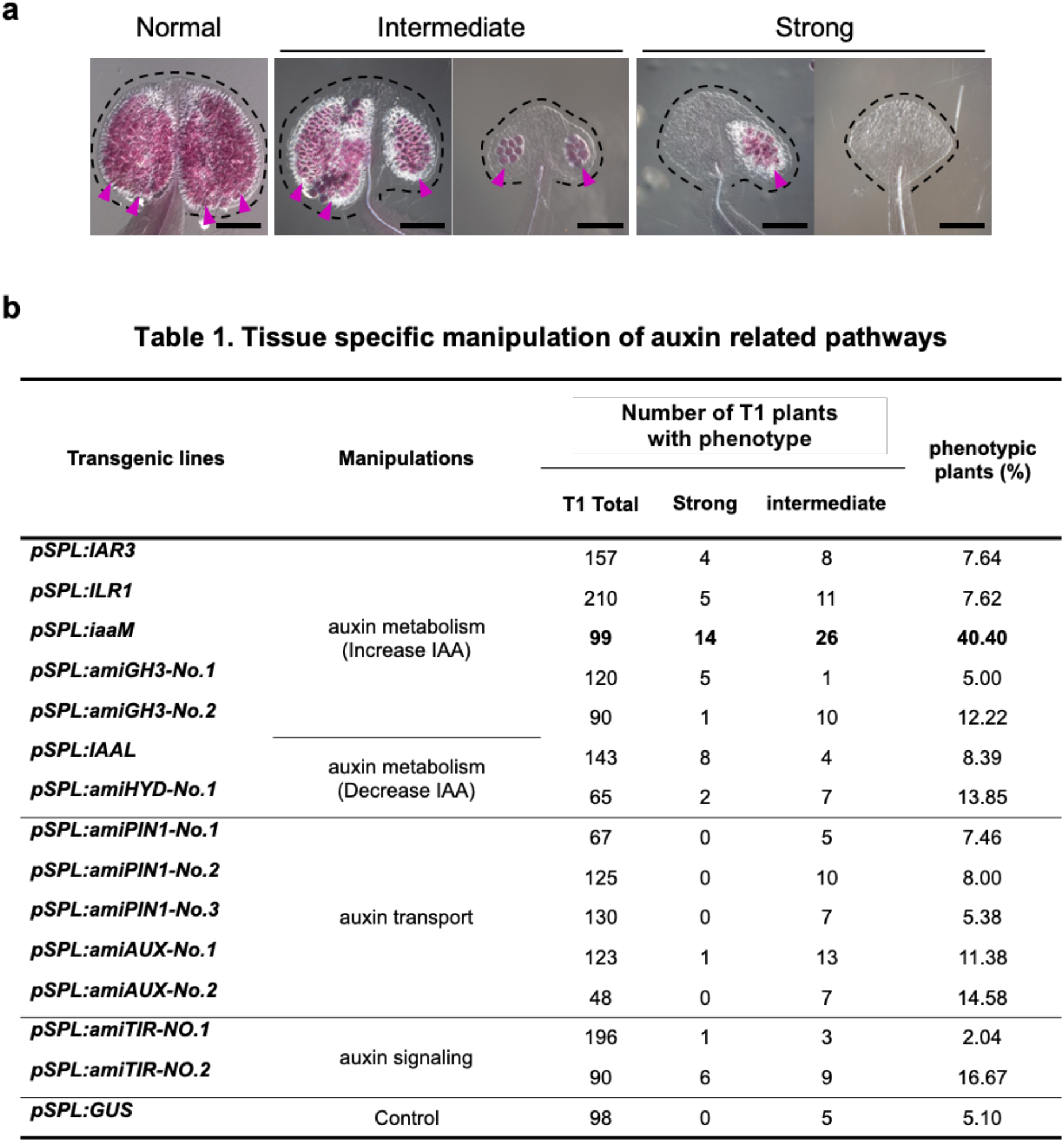
Large-scale testing the function of auxin in early anthers using transgenic strategy. **a**. Three categories of phenotype revealed by Alexander staining assay in transgenic plants. Normal anthers: 4 locules with pollens; intermediate phenotype: 2 or 3 locules with pollens; strong phenotype: no or only 1 locule with pollens. The shape of the anther was indicated with dotted lines. Magenta arrowheads indicate the locules with pollens (stained to red). **b**. Quantification of phenotypes in T_1_ transgenic plants according to the criteria shown in (a). Increasing IAA by expressing *pSPL:iaaM* caused the most severe sterility among all different perturbation conditions. Scale bars represent 100 μm.

**Figure S4.**
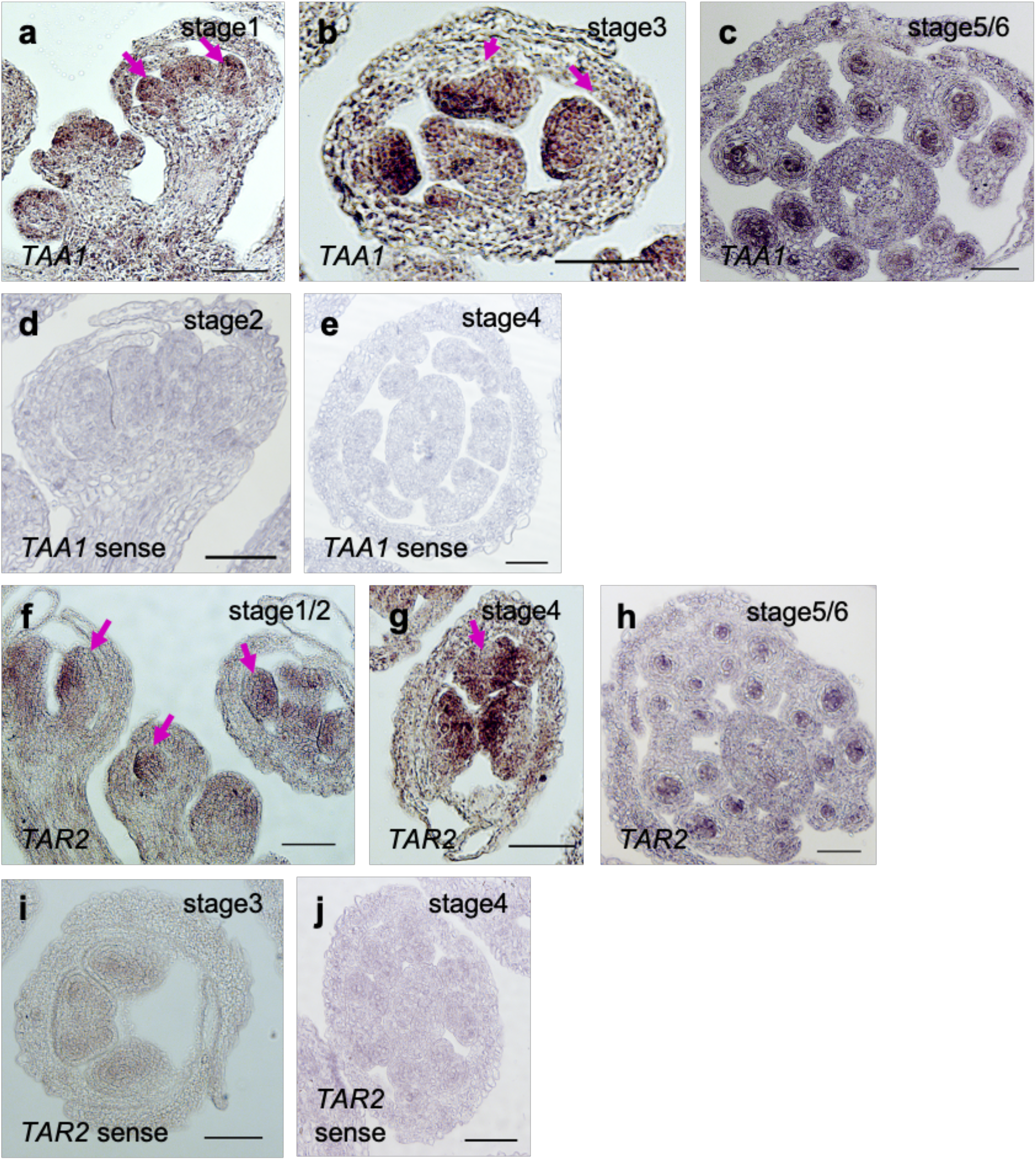
Expression pattern of *TAA1* and *TAR2* in Col anthers by *in situ* hybridization. **a-e**, TAA1 expression patterns in early anthers revealed by *in situ* hybridization with antisense (a-c) and antisense probe (d-e). **f-j**, TAR2 expression in early anthers by *in situ* hybridization with antisense (f-h) and antisense probe (i-j). Experiments were repeated 3 times and similar results were obtained. Arrows indicate anthers. Scale bars represent 50 μm.

**Figure S5.**
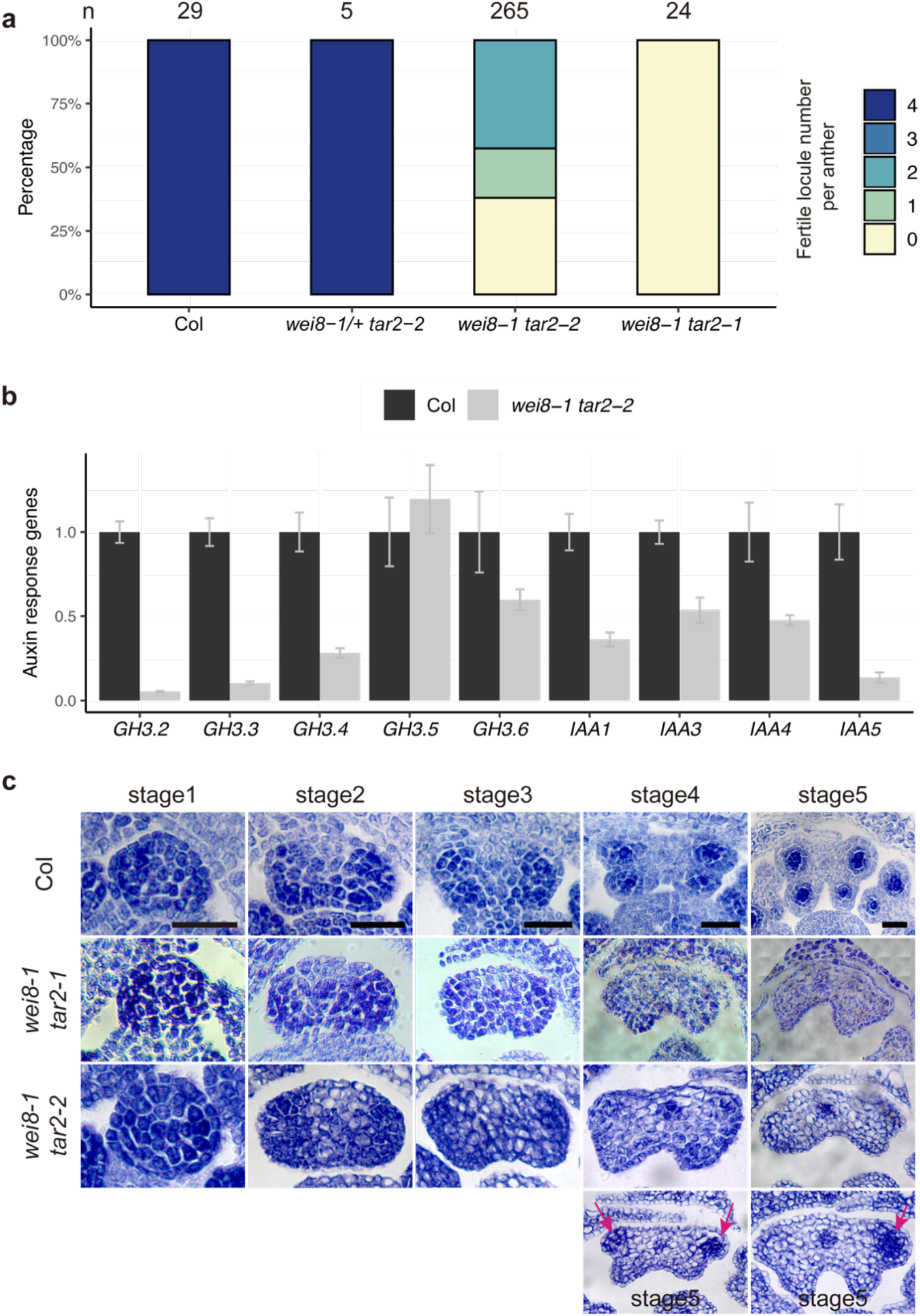
Phenotypic analysis of *wei8 tar2*. **a**. Quantification of anther fertilities in Col, *wei8-1/+ tar2-2/+*, *wei8-1 tar2-2*, and *wei8-1 tar2-1* by Alexander staining. Number of anthers (*n*) is shown above each individual bar. **b**. Expression of auxin responsible genes in Col and *wei8-1tar2-2* flowers by qRT-PCR. Bars indicate the mean, and the error bars indicate standard error (*n* = 3 technical repeats). Three independent experiments yielded similar results. **c**. Early anther development in Col, *wei8-1 tar2-1*, *wei8-1 tar2-2* from stage1 to stage 5 by cross sections. In some of *wei8-1 tar2-2* abaxial locules, GCs could be differentiated (magenta arrows). Scale bars represent 25 μm.

**Figure S6.**
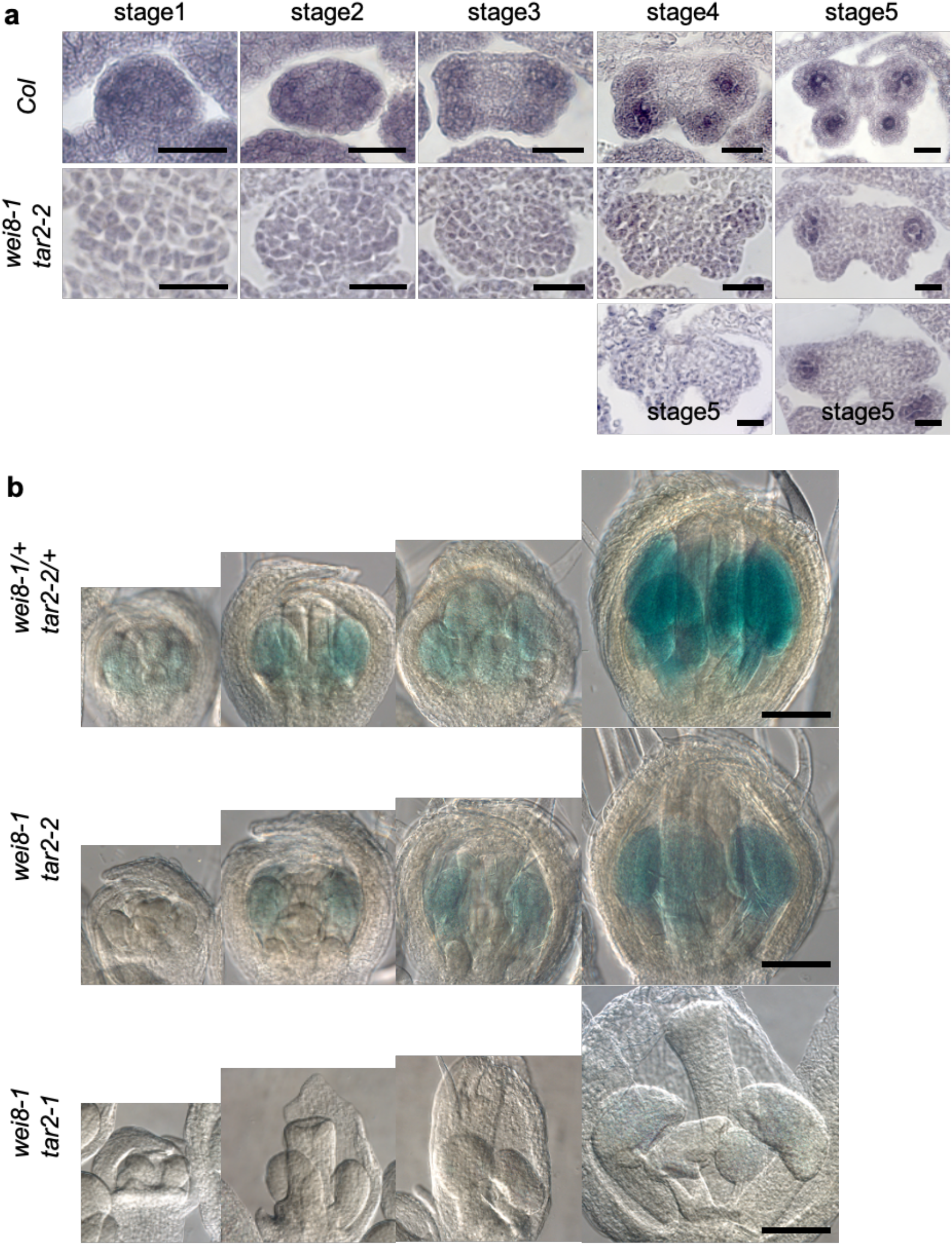
*SPL* transcript level in *wei8 tar2*. **a**. *In situ* of *SPL* in cross sections of Col and *wei8-1 tar2-2* anthers. **b**. The *SPL* promoter activity shown by *pSPL:GUS* signals in Col and *taa1 tar2*. Scale bars represent 25 μm in (a), and 1 mm in (b).

**Figure S7.**
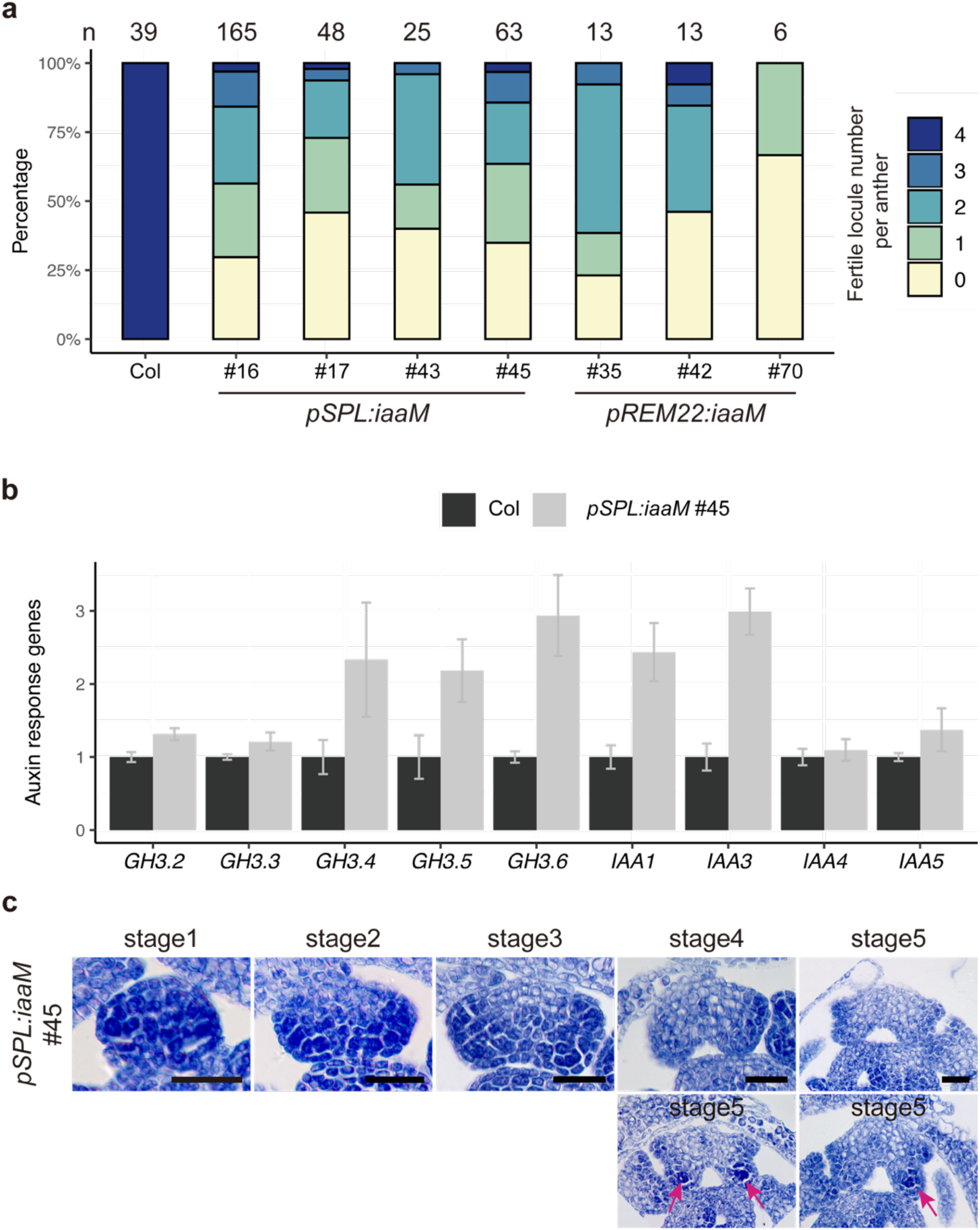
Increased auxin biosynthesis impairs anther development. **a**. Quantification of anther fertilities in Col, *pSPL:iaaM*, *pREM22:iaaM* by Alexander staining assay. Numbers above each bar indicate the number of observed anthers (n). **b**. Expression of auxin-responsive genes in Col and *pSPL:iaaM* flowers by qRT-PCR. Bars indicates the mean, and the error bars indicates standard error (*n* = 3 technical repeats). Three independent experiments yielded similar results. **c**. Early anther development in Col, *pSPL:iaaM* by section. Arrows indicate locules that cells are differentiated. Scale bars represent 25 μm.

**Table S1.**
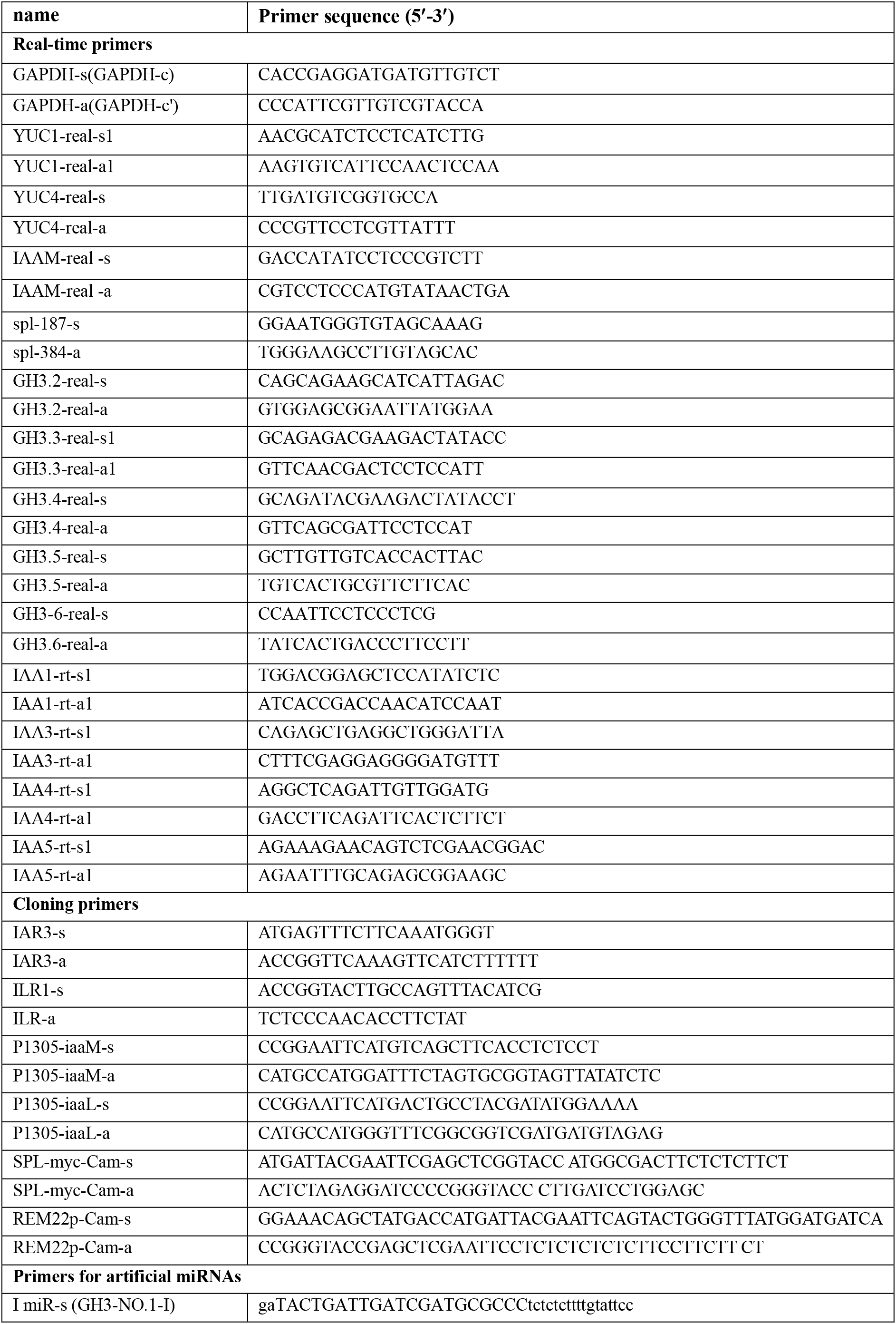
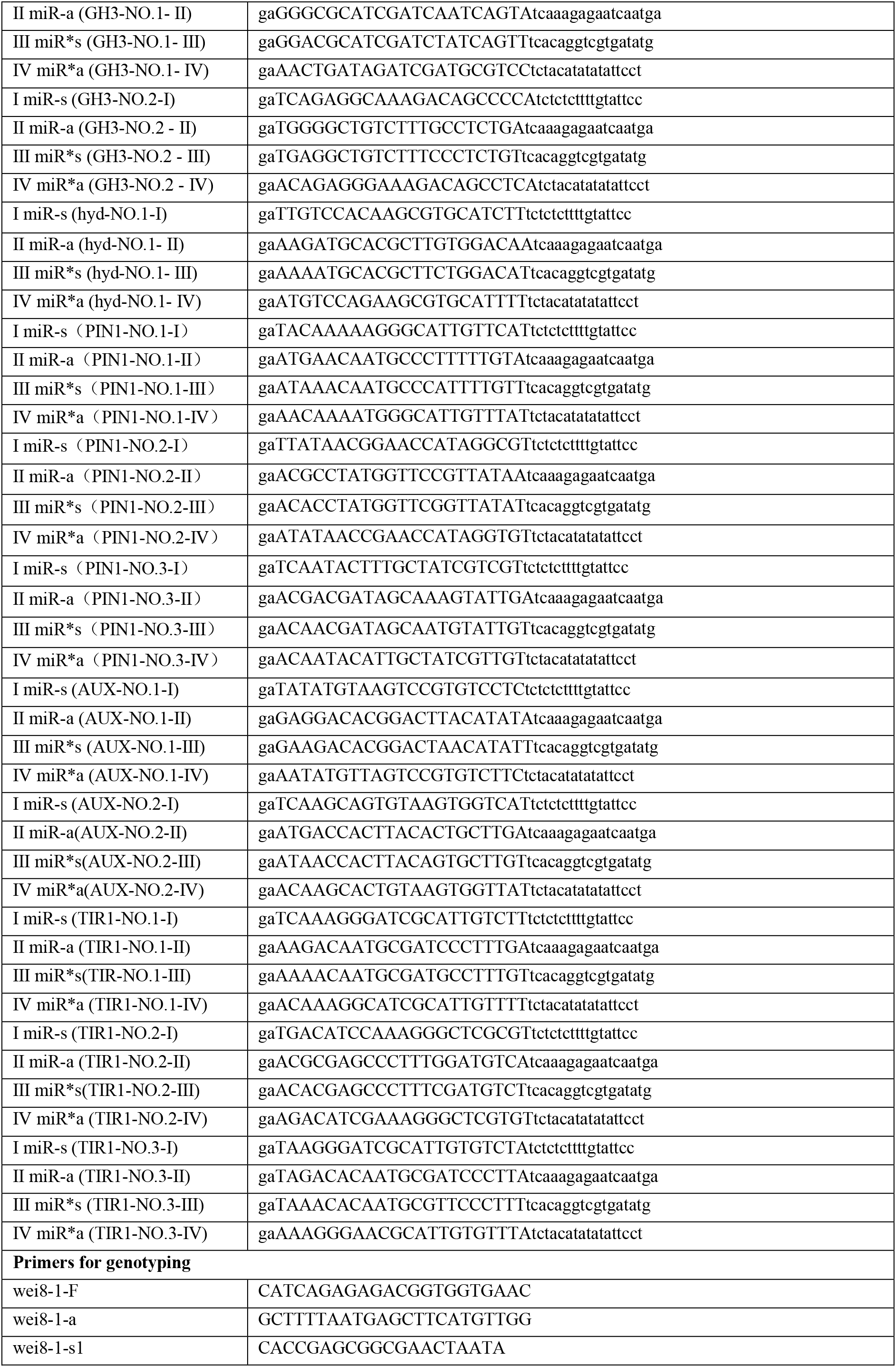
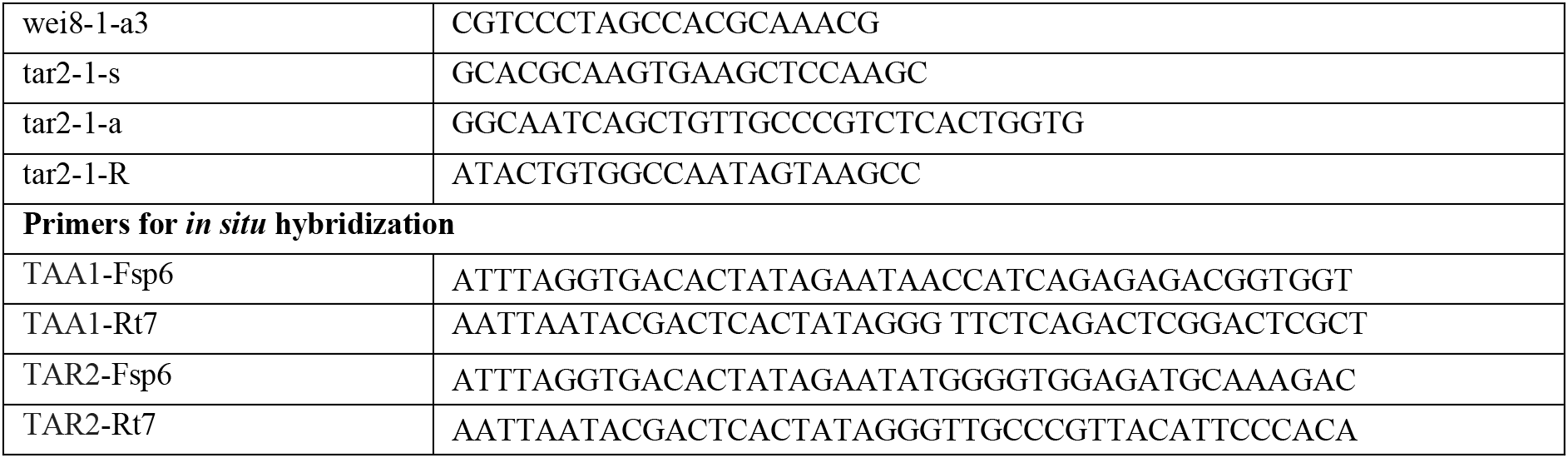
Primers used in this study.

